# A temporal map of maternal immune activation-induced changes reveals a shift in neurodevelopmental timing and perturbed cortical development in mice

**DOI:** 10.1101/2020.06.13.150359

**Authors:** Cesar P. Canales, Myka L. Estes, Karol Cichewicz, Kartik Angara, John Paul Aboubechara, Scott Cameron, Kathryn Prendergast, Linda Su-Feher, Iva Zdilar, Ellie J. Kreun, Emma C. Connolly, Jin M. Seo, Jack B. Goon, Kathleen Farrelly, Tyler Stradleigh, Deborah van der List, Lori Haapanen, Judy Van de Water, Daniel Vogt, A. Kimberley McAllister, Alex S. Nord

## Abstract

**Background:** Environmental insults that activate the maternal immune system are potent primers of developmental neuropathology and maternal immune activation (MIA) has emerged as a risk factor for neurodevelopmental and psychiatric disorders. Animal models of MIA provide an opportunity to identify molecular pathways that initiate disease processes and lead to neuropathology and behavioral deficits in offspring. MIA-induced behaviors are accompanied by anatomical and neurochemical alterations in adult offspring that parallel those seen in affected human populations.

**Methods:** We performed transcriptional profiling and neuroanatomical characterization in a time course across mouse embryonic cortical development, following MIA via single injection of the viral mimic polyinosinic:polycytidylic acid (polyI:C) at E12.5. Transcriptional changes identified in the cortex of MIA offspring at E17.5 were validated and mapped to cortical neuroanatomy and cell types via protein analysis and immunohistochemistry.

**Results:** MIA induced strong transcriptomic signatures, including induction of genes associated with hypoxia, immune signaling, and angiogenesis. The acute response identified 6h after the MIA insult was followed by changes in proliferation, neuronal and glial differentiation, and cortical lamination that emerged at E14.5 and peaked at E17.5. Decreased numbers of proliferative cell types in germinal zones and alterations in neuronal and glial cell types across cortical lamina were identified in the MIA-exposed cortex.

**Conclusions:** MIA-induced transcriptomic signatures in fetal offspring overlap significantly with perturbations identified in neurodevelopmental disorders (NDDs), and provide novel insights into alterations in molecular and developmental timing processes linking MIA and neuropathology, potentially revealing new targets for development of novel approaches for earlier diagnosis and treatment of these disorders.

## Introduction

Epidemiological association between maternal infection and NDDs has been found for autism spectrum disorder (ASD), schizophrenia (SZ), bipolar disorder (BPD), anxiety, and major depressive disorder (MDD) (1–4). Indeed, maternal immune activation (MIA) itself is sufficient to produce NDD-relevant outcomes in offspring in animal models (5,6). Offspring from female mice exposed to the toll-like receptor 3 (TLR3) agonist and viral mimic polyinosinic:polycytidylic acid [poly(I:C)], and offspring in maternal immune activation models using the bacterial mimic lipopolysaccharide (LPS) (7–9), recapitulate neuropathologies and aberrant behaviors seen in offspring born to flu-infected dams (10,11). MIA in mice produces behavioral and cognitive abnormalities with relevance across multiple NDD diagnostic boundaries (12–14).

MIA-induced outcomes in adult offspring have been well characterized, however, mechanisms of initiation and progression of NDD-related brain pathology remain poorly defined. Poly(I:C) increases proinflammatory cytokines such as IL-1β, IL-6, CXCL1, and TNF-α in the maternal circulation (11,15–19). Downstream pathological changes in the brains of offspring have been reported across cell types, from neurons to microglia, and these changes have been linked to perturbations in proliferation, migration, and maturation in the developing brain (20–24). Recent studies have identified acute transcriptomic changes in fetal brain in mice and rats at single time-points (3-4h) following MIA, with significant overlap between transcriptome changes in the cortices of MIA-exposed offspring and those altered in the brains of children with ASD (25). Although MIA must cause a dynamic sequence of transcriptional changes in the fetal brain following acute exposure, no integrated picture of MIA-associated transcriptomic pathology exists that maps changes spanning from the initial acute response to MIA through birth. Here, we combined time course transcriptomics with neuroanatomy to map changes in prenatal mouse cortical development following MIA induced by poly(I:C) injection at E12.5.

## Methods and Materials

### Animal Care and Use

All studies were conducted in compliance with NIH guidelines and approved protocols from the University of California Davis Animal Care and Use Committee. C57BL/6N females (Charles River, Kingston, NY), were bred in house and maintained on a 12:12 h light dark cycle at 20 ± 1 °C, food and water available *ad libitum*. Mice utilized in this study do harbor Segmented Filamentous Bacteria (SFB) (26). Males and females embryos were analyzed across experiments, sex was determined as previously described (27).

### Maternal Immune Activation

Assessment of females baseline immunoreactivity before pregnancy and maternal immune activation were performed as previously described (28). Since female mice display a wide range of baseline immunoreactivity to poly(I:C) that dictates susceptibility and resilience of offspring from later pregnancies to MIA, virgin females in the lowest 25^th^ percentile of immune responsiveness to poly(I:C) (IL-6 serum levels) were excluded from the study to reduce variability and ensure sufficient immune responsiveness of all included dams. E12.5 pregnancies were determined by visualizing vaginal seminal plug (noted as E0.5) and by body weight increase. IL-6 serum levels were measured at 2.5 h post-injection. Temperature, weight, and sickness behavior were tracked as previously described (28).

### Gross anatomy, protein validation, and lamination image analyses

Histology was performed in triplicates on brains representing at least two litters, selected randomly without taking morphological criteria into account. Morphological parameters were measured using FIJI ImageJ (NIH), by an experimenter blinded to group and sex. Brain regions were identified based on anatomical landmarks as previously described (29). No data points were excluded. Immunohistochemistry staining and immunoblotting were performed following conventional protocols with techniques and antibodies described in supplementary material.

### Transcriptomics

Dissections were performed at E12.5 + 6hrs, E14.5, E17.5, and P0. Pallial and subpallial structures were included in tissues dissected at E12.5. Later time points did not include subpallium. Dissections included both hemispheres from males and females, with exact numbers and sample details reported in **Supplementary Table 1** Total RNA was isolated using Ambion RNAqueous Total RNA Isolation Kit and assayed using Agilent RNA 6000 Nano Bioanalyzer kit/instrument. Stranded mRNA libraries were prepared using TruSeq Stranded mRNA kits. 8–12 samples per lane were pooled and sequenced on an Illumina HiSeq 4000 instrument using a single-end 50-bp protocol. Reads were aligned to mouse genome (mm9) using STAR (version 2.4.2a) (30) and gene counts produced using featureCounts (31) and mm9 knownGenes. Quality assayed using FastQC (32) and RSeQC (33), and samples that exhibited 3′ bias or poor exon distribution were discarded. Detailed methods including parameters for differential expression, gene set enrichment, and WGCNA reported in supplementary material.

### Data Analysis

For all experiments, data was collected from at least two (usually three, and sometimes more, as indicated in each analysis) experiments and is presented as the mean ± SEM. Protein validation data were analyzed by Student’s *t*-test or one-way ANOVA, followed where appropriate by Tukey’s honestly significant difference *post hoc* test (Graphpad Prism v.7). Significance was defined as: *p <0.05, **p <0.01, ***p <0.001, ****p <0.0001. Samples for RNA-seq were randomly collected across litters and processed blind to experimental condition. For differential gene expression analysis using edgeR, differences were considered statistically significant at high stringency for FDR < 0.05, or with reduced stringency for *P* values < 0.05.

## Results

Induction of MIA via poly(I:C) injection at E12.5 produces relevant phenotypes in mice (11,20,34) and the specific implementation of this model was recently validated in our hands to produce sufficient levels of MIA to mimic viral infection and cause aberrant behavioral outcomes in offspring (28). The dosage of 30 mg/ml poly(I:C) elicited substantial elevations in maternal serum IL-6 (on average 2,200 pg/ml) 4 hours following injection **(Supplementary Fig. 1)**. Paired transcriptomic and neuroanatomical analyses were used to map changes in the fetal brain following MIA in a poly(I:C) mouse model (**Fig 1a**). Pregnant mice were injected with saline or poly(I:C) at E12.5, and fetal forebrain (pallial and subpallial structures, E12.5 + 6hrs) or cerebral cortices (E14.5, E17.5, and birth (P0)) were microdissected. RNA-seq datasets were generated from male and female embryos from 28 independent litters across control and MIA groups, typically with 1-3 embryos represented per independent litter.

**Figure 1:**
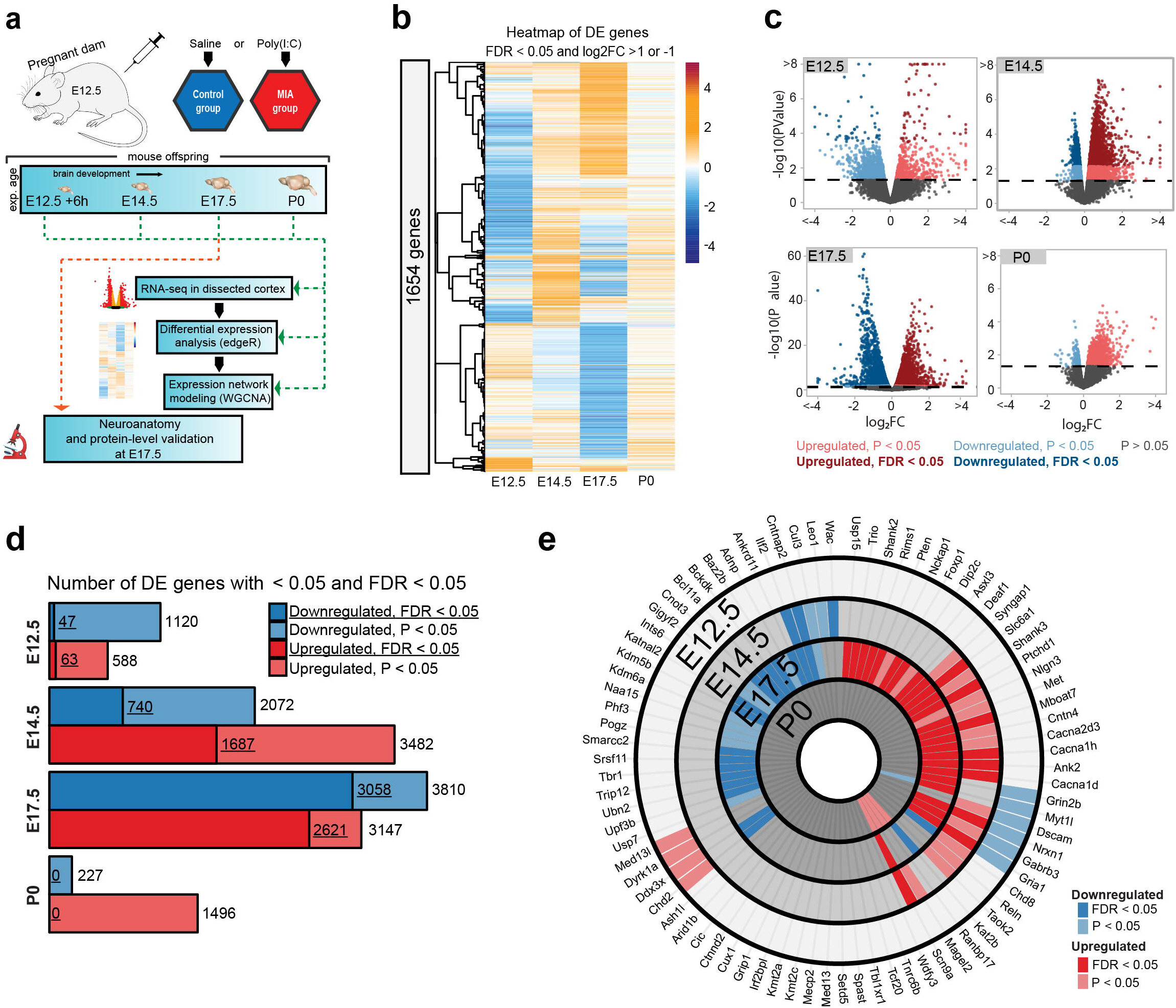
Differential expression across embryonic brain development following maternal immune activation via poly(I:C) injection at E12.5. **(a)** Schematic representation of MIA model and experimental pipeline. E: embryonic day, P: postnatal day, IHC: immunohistochemistry, WB: Western blot. Poly(I:C) was injected at E12.5. Samples for RNA-seq were collected at E12.5 + 6h, E14.5, E17.5, and at birth (P0). IHC and WB analysis were conducted on a separate animal cohort at E17.5 and P0. **(b)** Heatmap representing relative gene expression changes between control and MIA samples across neurodevelopment. X-axis, time point. Y-axis, hierarchically clustered DE genes (FDR < 0.05 and log_2_ fold change (log_2_FC) > 1 or < −1). **(c)** Volcano plots of DE effect size and significance. Colors represent directionality and statistical significance. **(d)** Numbers of upregulated and downregulated DE genes at FDR < 0.05 and p <0.05 thresholds. **(e)** Overlap of DE genes with 82 SFARI autism-associated mouse gene orthologs. Concentric circles represent developmental timepoints. Light red, upregulated DE genes (p <0.05); dark red, upregulated DE genes (FDR < 0.05); light blue, downregulated DE genes (p <0.05); dark blue, downregulated DE genes (FDR < 0.05). SFARI gene orthologs are statistically enriched among E17.5 DE genes for genes passing FDR < 0.05 (P = 3.9e-04, hypergeometric test) and p <0.05 (P = 2.8e-06, hypergeometric test).

Comparison of MIA and control samples identified robust differential expression (DE) signatures stratified by developmental stage (**Fig. 1b**, **Supplementary Fig. 2a, b)**. For each time point, DE genes were defined using a stringent false discovery rate (FDR) < 0.05 threshold and a more inclusive p <0.05 threshold (**Fig. 1b, c**, **Supplementary Table 2-6)**. Sex-stratified analysis did not indicate strong DE differences between male and female cortices following MIA, though DE effect sizes appeared stronger in females at E14.5 and E17.5, leading to a larger number of DE genes identified independently in females compared to males (**Supplementary Fig. 3a – d, Supplementary Table 7-16**). Overall, sex-stratified differences in DE were generally subtle, as demonstrated by high DE correlation and shared DE gene sets between sexes, with consistent findings across sexes among key DE genes (**Supplementary Fig. 4a-c**). Samples from both male and female offspring were used for overall DE analysis models, with sex included as a covariate.

For the DE gene set passing the stringent FDR < 0.05 threshold, there were varying numbers of DE genes across timepoints **(Fig. 1d)**. The strongest transcriptional signature was observed at E17.5 (2621 up and 3058 down), suggesting dramatic impact of MIA on cortical development at this time point. P0 represented the subtlest transcriptomic signature, with no genes passing the stringent FDR < 0.05 threshold, and with the P <0.05 DE genes showing a strong upregulation bias. We tested for overlap between DE RNA-seq genes and high confidence autism-associated genes in the Simons Foundation Autism Research Initiative (SFARI) gene database (35). The 82 mouse orthologs of SFARI ASD genes that were expressed at measurable levels were significantly enriched among E17.5 DE genes, with 25 upregulated and 19 downregulated DE genes at FDR < 0.05 (P = 3.9e-04, hypergeometric test), and 28 upregulated and 28 downregulated DE genes at p <0.05 (P = 2.8e-06, hypergeometric test) **(Fig 1e**, **Supplementary Table 17-18)**. This enrichment during peak transcriptomic dysregulation suggests involvement of ASD-relevant pathways in MIA etiology here. Detailed results of DE analysis are reported in **Supplemental Tables 2 – 16** and can be visualized using our interactive online browser (https://nordlab.shinyapps.io/mia_browser/).

We next sought to capture MIA-induced gene regulatory changes at the systems and network level across embryonic cortex development using weighted gene co-expression network analysis (WGCNA) (36,37). In this approach, genes with correlated expression patterns are assigned into modules, enabling identification of gene sets with shared expression and function in neurodevelopment. WGCNA co-expression modules were arbitrarily named after colors, with Grey reserved for genes with no strongly correlated expression patterns **(Fig. 2a, b)**. Our initial analysis identified 10 modules **(Supplementary figure 5a)**, of which correlated modules were combined, and the 6 final modules captured dynamic trajectories of neurodevelopmental gene expression **(Fig. 2b)**. The Blue and Turquoise modules represented genes that increased or decreased, respectively, in expression during neurodevelopment (Blue: P = 9e-34; Turquoise: P = 1e-29, **Supplementary Fig. 5b**). The other modules showed more complex patterns of expression. Each WGCNA module included DE genes, with differences in what time point had the most extensive DE, indicating module-specific timing of perturbation associated with MIA **(Supplementary Fig. 5c)**. DE genes were enriched for module-specific GO terms, with GO enrichment findings similar using all module genes and using an alternative rank-based gene set enrichment that is not dependent on FDR or P-value (38,39) **(Fig 2c**, **Supplementary Table 19; Supplementary Fig. 6a-h, Supplementary Table 20)**.

**Figure 2:**
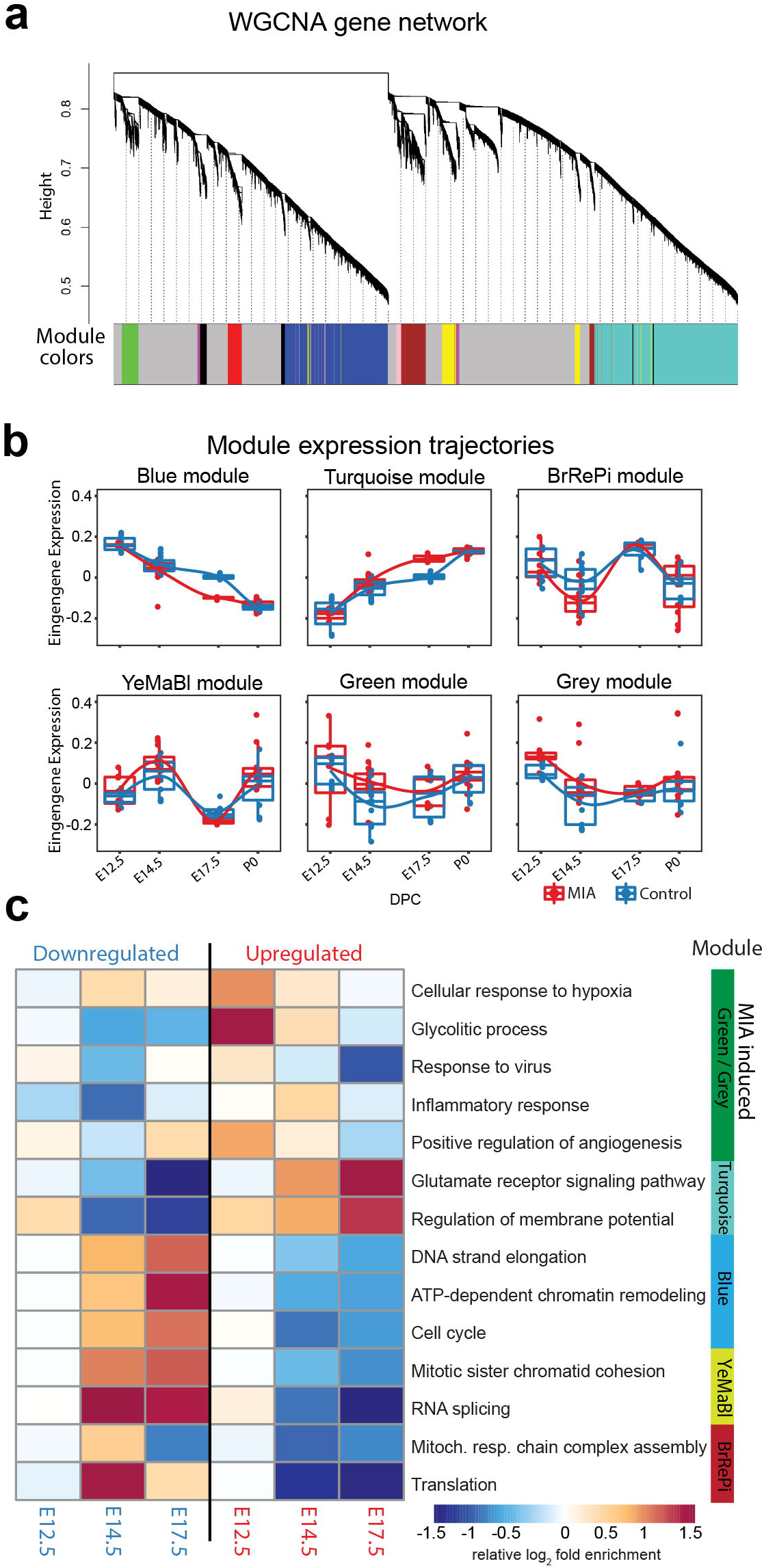
Weighted gene co-expression network analysis (WGCNA) reveal discrete gene networks perturbed by MIA across embryonic cortical neurodevelopment. **(a)** WGCNA cluster dendrogram of time series samples grouping genes into co-expression modules. **(b)** Trajectories of module eigengene expression across neurodevelopment for MIA and control samples. **(c)** Heatmap of representative gene ontology biological processes (GO BP) across developmental timepoints enriched in co-expression modules. Y-axis represents enriched GO BP terms, among DE genes passing FDR < 0.05, marked for their module association (p <0.05, Fisher’s exact test, in at least one timepoint). Heatmap color scale represents relative fold enrichment. P0 not shown due to insufficient DE gene numbers and absence of GO BP terms passing enrichment criteria.

The WGCNA-resolved modules enabled mapping of DE signatures to neurodevelopmental processes that were acutely induced by MIA. There was significant association between genes in Grey (P = 0.007) and Green (P = 0.03) modules and MIA treatment, and marginal significance with MIA for the YeMaBl (P = 0.05) module **(Supplementary Fig. 5b)**. The acute, initiating signaling pathways found six hours after poly(I:C) injection captured by the Green module and by a subset of genes within the Grey module, show a strong signature of upregulated DE genes from E12.5 to E14.5 **(Fig 3a).** Analysis of the GO BP enrichment of upregulated DE genes in these modules identified activation of immune-related pathways, including defense response to virus, as well as angiogenesis and VEGFA signaling **(Fig. 2c**, **3b**, **Supplementary Fig. 6a, c)**. We tested for protein-protein association networks among the *acute E12.5 DE (FDR < 0.05) genes using STRING (40), and found significantly more* interactions than expected by chance (observed edges = 161, expected edges = 46, enrichment = 3.5, STRING P <1.0e-16) (**Fig. 3c**). This intersecting protein network was particularly associated with metabolism, hypoxia, and oxidative stress **(Fig. 3b)**. The network includes a highly interacting core gene set with hub nodes, such as *Vegfa*, that have many spokes and may be signaling factors that direct the MIA response **(Fig. 3b, c)**. Developmental expression profiles for DE genes that were associated with this interaction network, are shown in **Fig. 3d**, and stratified by sex in **Supplementary Fig. 7a**. These signatures capture a complex but coherent transcriptional responses involving hypoxia, immune, metabolic, and angiogenesis pathways that are strongly and transiently induced in fetal brain within 6 hours following mid-gestational MIA.

**Figure 3.**
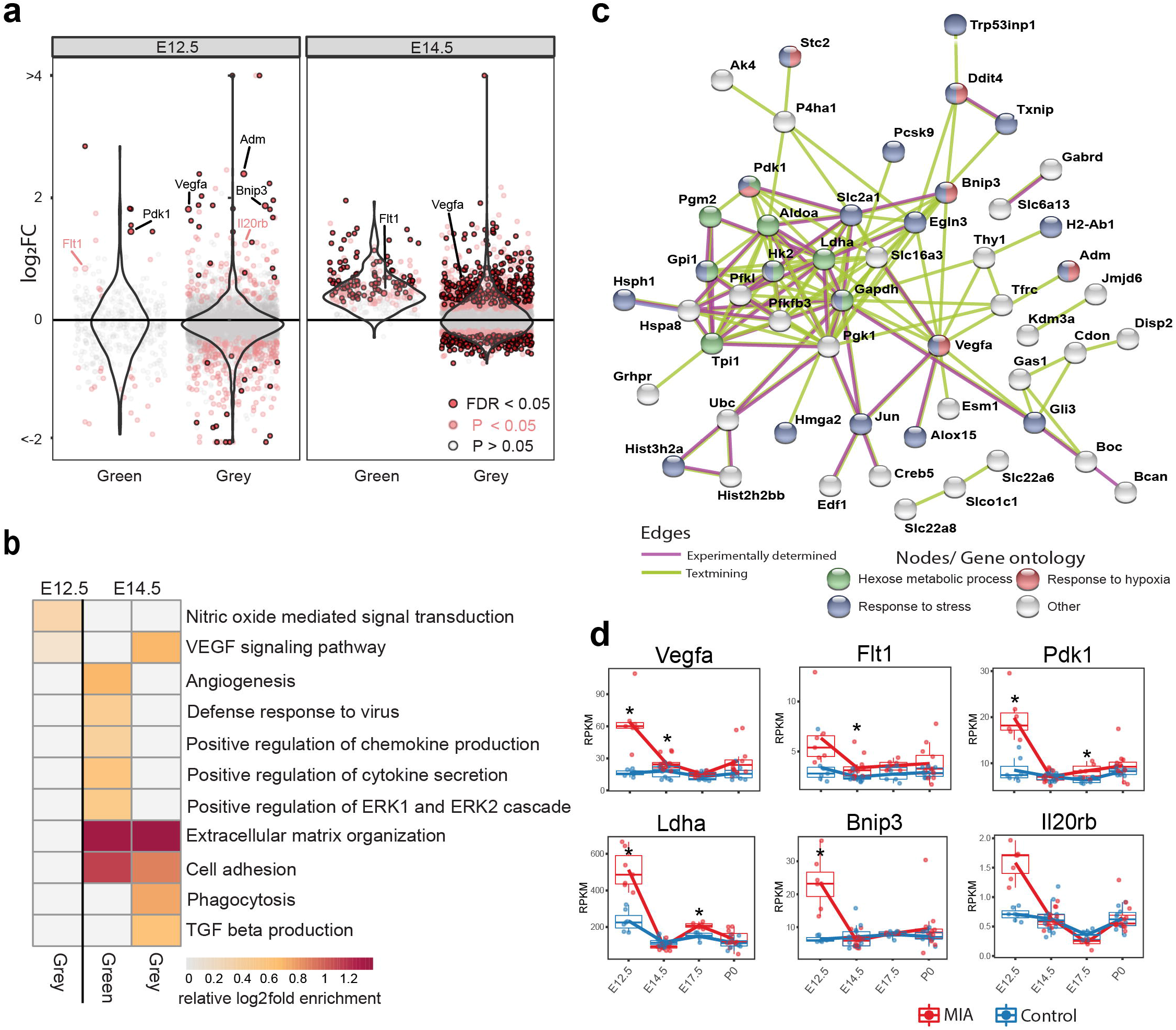
Green and Grey co-expression modules capture acute transcriptional response at six hours and 2 days following MIA exposure. **(a)** Log_2_ fold changes of Green and Grey module gene expression between control and MIA animals. Violin plots visualize distribution of log_2_ fold changes. Genes, which expression trajectories are plotted in (d) are labeled. **(b)** Gene set enrichment analysis of GO BP terms significantly enriched among FDR < 0.05 genes at E12.5 and E14.5 in Green and Grey modules identified upregulation of angiogenesis and immune pathways. Representative GO BP terms with p <0.05; grey color represents enrichment with P > 0.05 (Fisher’s exact test). **(c)** STRING protein–protein interaction network of E12.5 DE genes (FDR < 0.05) colored by annotation to GO biological process terms. **(d)** RPKM expression plots of genes labeled in (a). Stars represent FDR < 0.05.

The most pervasive WGCNA-resolved DE signatures following MIA impacted a large proportion of genes in the Blue and Turquoise modules by E14.5 and peaked at E17.5 **(Fig. 1c)**. GO terms related to proliferation and cell cycle were enriched in the Blue module among downregulated DE genes, and terms related to neuronal differentiation and synapses were enriched in the Turquoise module among upregulated DE genes **(Fig. 2c)**. Considering the changes in proliferative and lamination markers associated with Blue and Turquoise genes **(Fig. 4a**, **Supplementary Fig. 7b**), we sought to validate such changes at the protein level in independent samples. Four out of 5 DE genes tested validated with concordant changes at the protein level as determined by Western blot (WB) analysis at E17.5, showing either statistical significance or trends in the same direction as observed in our transcriptomic analysis **(Fig. 4b**, **Supplementary Fig. 8).** The DE signatures associated with Blue and Turquoise modules is evidenced by separation between MIA and saline samples in PCA plots at E14.5 and E17.5, with E17.5 MIA samples clustered closer to the P0 samples than saline age-matched counterparts **(Supplementary Fig. 2)**. To test for MIA-associated perturbation to temporal full transcriptome expression patterns, we generated a linear regression model using RNA-seq data principal components 1-5 to model age in saline samples. We then used this model to predict age in MIA samples **(Fig. 4c)**. E17.5 MIA samples exhibited older predicted age values relative to E17.5 saline controls (P = 4.5e-06, T-test), and E14.5 MIA samples showed a similar trend (P = 0.07, T-test). These findings suggest that mid-gestational MIA at E12.5 perturbs the timing of major downstream neurodevelopmental processes at E14.5 and E17.5 in embryonic cerebral cortex.

**Figure 4.**
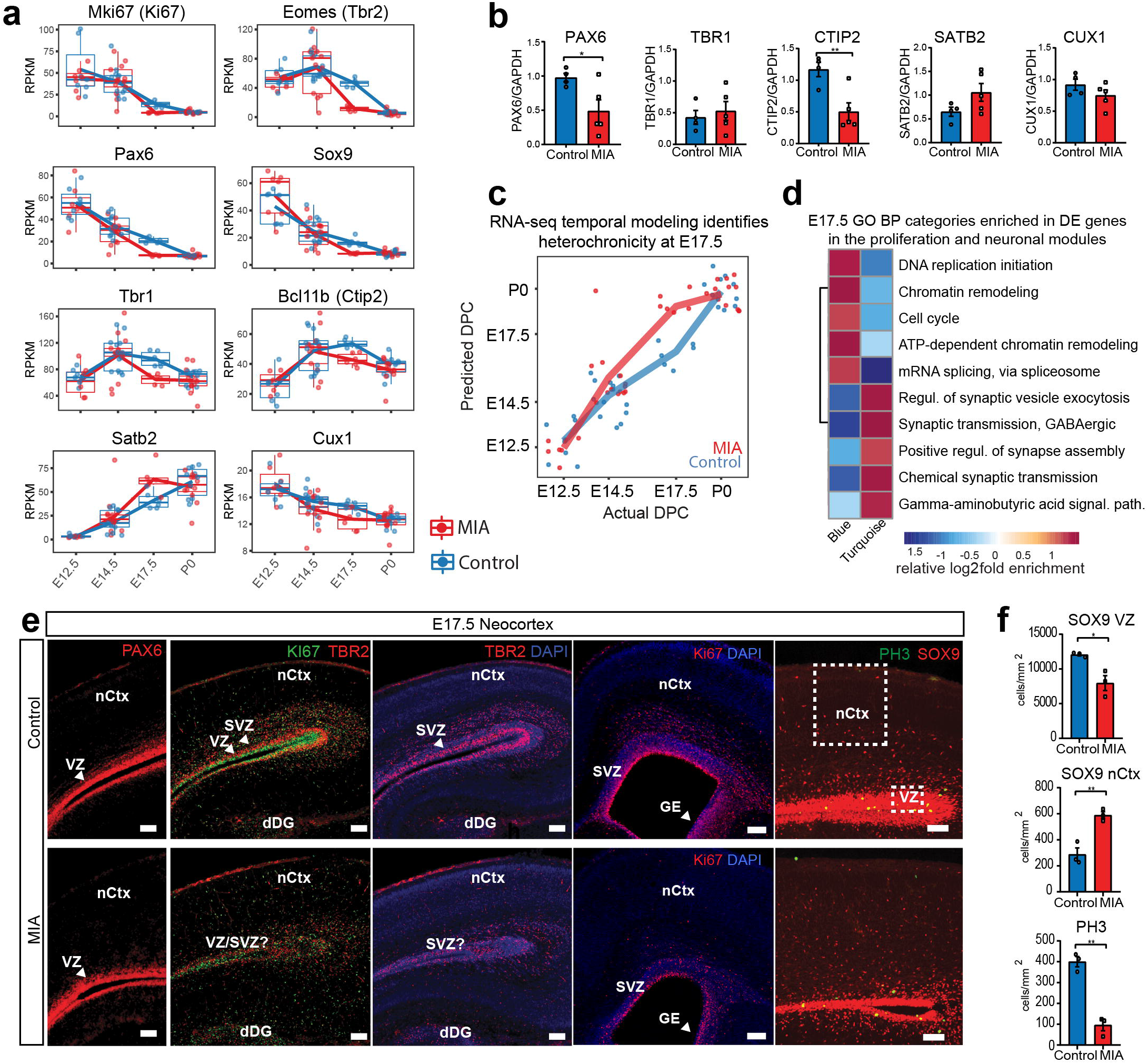
MIA profoundly alters cortical proliferation and lamination dynamics in the fetal offspring five days after poly(I:C) injection. **(a)** RPKM trajectories of proliferative and cortical lamination markers validated at protein level. **(b)** Quantified protein levels from WB analysis for the DE genes at E17.5 relative to GAPDH protein expression, (n = 4 control, n = 5 MIA; PAX6 P = 0.039, TBR1 P = 0.615, CTIP2 P = 0.0090, SATB2 P = 0.1160, CUX1 P = 0.2112; two tailed Student’s t-test, individual blots in supplementary data **(Supplementary Fig. 10 a-c, h, i)**. **(c)** Temporal modeling of the RNA-seq data suggests alteration of the neurodevelopmental program in MIA animals at E17.5 (P = 4.5e-06, Student’s t-test) and similar trend at E14.5 (P = 0.07, Student’s t-test). Actual age (X-axis) vs predicted age (Y-axis) calculated by the linear model. Control samples were used for training the model. Lines connect average values, points depict individual samples and are jittered along the X-axis. **(d)** GO BP categories enriched in DE genes in the proliferation (Blue) and neuronal (Turquoise) WGCNA modules. **(e)** Coronal brain sections from E17.5 saline (control brains, n=4 from 2 independent litters) and MIA animals (n=6 from 2 individual litters) were immunostained to detect: progenitor cells (PAX6, SOX9), proliferative cells (KI67, PH3), and intermediate progenitors (TBR2) (nCtx: neocortex, VZ: ventricular zone, SVZ: sub-ventricular zone, dDG: developing dentate gyrus). Staining for all cell populations is qualitatively decreased in the cortices from MIA offspring. Representative images are shown. Boxes in top right panel indicate regions of the VZ and upper layers of the nCtx utilized for SOX9 cell counts. Scale bars = 100 μm. **(f)** SOX9 positive cell density quantifications in the VZ as well as in the upper layers of the nCtx show reduced and ectopic SOX9+ cells in the MIA group (n = 3 per condition, SOX9 VZ *P = 0.0198, SOX9 nCtx **P = 0.0058, two tailed Student’s t-test) and PH3 cell density quantification along the entire length of the VZ confirms reduction (n = 3 per condition, **P = 0.0012, two tailed Student’s t-test).

To characterize the neuroanatomical specificity of Blue and Turquoise dysregulation transcriptional signatures **(Fig. 4d)**, we performed a static immunohistological comparison of late neurogenesis, cortical lamination and cell specification at E17.5 in an independent MIA cohort. E17.5 represents the end of the peak of neurogenesis during cortical development, before the neuronal-glial transition occurs around E18.5. At this stage, it is expected that progenitor markers, such as PAX6 and SOX9, will identify a small population of radial glia that are maintained and retain the ability to produce neurons, and also subpopulations of proliferative cells commencing gliogenesis (41). PAX6+ and SOX9+ cell distribution patterns showed that progenitors localized correctly around the cortical ventricular and subventricular zones in coronal mid-telencephalic sections; however, both PAX6+ and SOX9+ populations were significantly reduced in MIA brains compared to controls **(Fig. 4e)**. The PAX6 finding is consistent with our WB analysis (PAX6, P = 0.039 at E17.5, two tailed Student’s t-test) **(Fig. 4b**, **Supplementary Fig. 8a)**. While MIA brains showed a decrease in SOX9+ cells in the ventricular zone (VZ), increased SOX9+ cells above the VZ in the neocortex were observed (SOX9 VZ, P = 0.0198; nCtx, P=0.0058, two tailed Student’s t-test) (**Fig. 4f**). PAX6 protein levels either resolved or were too subtle to be distinguishable by birth (P = 0.766, two tailed Student’s t-test) **(Fig. 4a**, **Supplementary Fig. 9b, Supplementary Fig. 8b-c, i).** We next performed IHC for KI67 and PH3, active cell cycle state markers, and Tbr2, a marker for post-mitotic intermediate progenitors. Analysis of dividing KI67+ and PH3+ cells showed reduced neurogenesis in the VZ along with a qualitative reduction in numbers and ectopic positioning of postmitotic intermediate progenitor cells in the MIA group (TBR2+). This reduction in Ki67+ cells was observed in several regions of the developing cortex, including zones defined as S1 and S1DZ (20,34), as well as the ganglionic eminences **(Fig. 4e)**. Quantification of the PH3+ cells confirmed the MIA-induced reduction in mitosis (PH3, P = 0.0012, two tailed Student’s t-test) **(Fig 4f)**. Thus, altered transcriptomic proliferative dynamics were linked to changes at the protein level and with decreased proliferative cellular populations at E17.5 following MIA.

There are conflicting results as to whether MIA alters cortical thickness and/or causes cortical dysplasia in young adult MIA offspring (11,24,42), with stereotypic dysplasia impacting lamination localized to the somatosensory cortex reported in one MIA model (20,34). A number of cortical layer markers were DE at E17.5 in RNA with validated changes at the protein level **(Fig. 4 a-b**, **Supplementary Fig. 7b)**. To examine cortical lamination, distribution of cortical layer-specific markers was analyzed via IHC in the same samples as above. CTIP2 and TBR1 staining revealed deep cortical layer-specific alterations in lamination, thickness, and cell density (**Fig. 5a-c)**. MIA brains appeared to be at a more complete stage of lamination at E17.5, showing cortical deep layers fully separated, as opposed to control animals that presented a CTIP2/TBR1 partial layer overlap at the same age **(Fig. 5a-b)**. The number of TBR1+ and CTIP2+ cells as well as their layer-specific thickness were also significantly reduced in the neocortex from MIA offspring (TBR1+ cells, P = 0.0138; CTIP2+ cells, P = 0.0119; TBR1 thickness, P = 0.075; CTIP2 thickness, 0.024; two tailed Student’s t-test). Thickness of more superficial cortical layers, represented by SATB2+ and CUX1+ cells, were significantly increased and unchanged, respectively (SATB2, P = 0.042; CUX1, P = 0.430, two tailed Student’s t-test) **(Fig. 5c-d)**. These findings are consistent with individual trajectories of DE genes and WB analyses **(Fig. 4a-b**, **Supplementary Fig. 8h)**. Finally, to assess cortical gross anatomy, we looked for MIA-associated dysplasic alterations across the brain, and measured the thickness of the developing somatosensory cortex at two anteroposterior regions at E17.5. We did not find obvious cortical dysplasia or gross alterations in brain anatomy in the analyzed rostral and caudal regions, including somatosensory cortex and S1DZ **(Fig. 5e-f)**.

**Figure 5:**
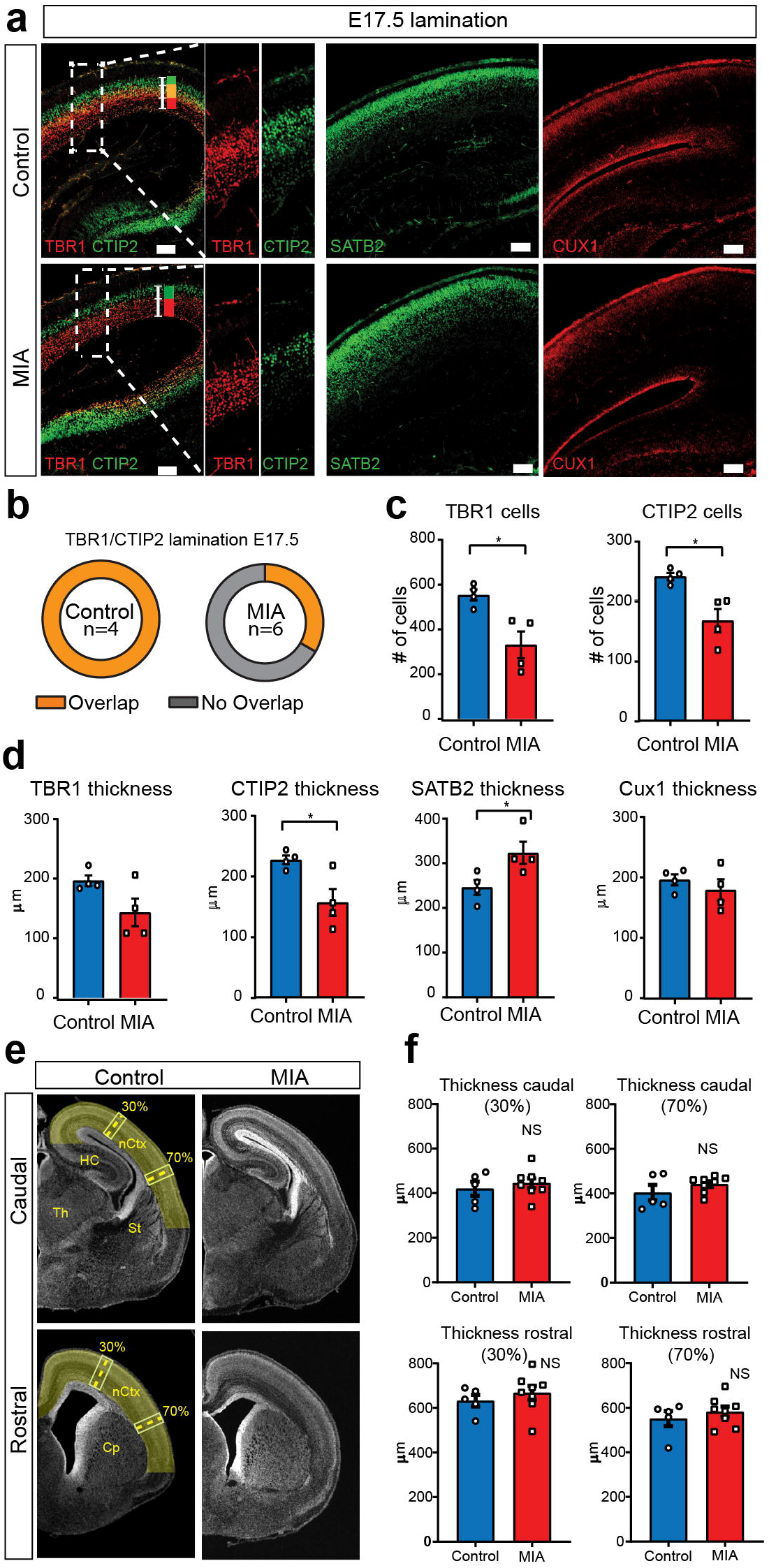
MIA impacts cortical lamination during development with subtle changes that do not alter cortical gross anatomy. **(a)** Coronal fetal brain sections from E17.5 saline (control) and MIA offspring brains were immunostained using antibodies to identify deep cortical lamination layer V, CTIP2 (green), and layer VI, TBR1 (red) and markers of postmitotic neurons, SATB2 and CUX1, in superficial cortical layers in brains derived from saline (control) or MIA injected mothers. Representative images are shown. CTIP2/TBR1 overlap, represented by yellow color, observed in control brains, compared to most MIA brains that presented fully laminated deep layers, with no overlap illustrated by adjacent green and red rectangles. Boxes indicate magnified areas used for quantification and CTIP2/TBR1 overlap analyses. Scale bars = 100 μm. **(b)** TBR1/CTIP2 lamination overlap occurrence. Brains from 3 MIA litters and 2 control litters were co-stained with TBR1 and CTIP2 cortical markers (as shown in a-b) and analyzed for lamination overlap occurrence (n=4 control, n=6 MIA). While all control brains showed lamination overlap between TBR1/CTIP2 layers, absence of TBR1/CTIP2 lamination overlap was observed in 4 out 6 analyzed MIA brain **(c)** TBR1+ and CTIP2+ cell counts and **(d)** layer-specific thickness analysis, based on the spread of the distribution of labeled cells, indicate decreases in both cortical markers in MIA brains. SATB2 and CUX1 layer-specific thickness analysis indicate expansion of the superficial layer delimited by SATB2+ cells in MIA brains (Cell counts: n = 4 brains per condition, TBR1 P = 0.0138; CTIP2 P = 0.0119. Layer-specific thickness analysis: n = 4 per condition, TBR1 P = 0.075; CTIP2 P = 0.0239; SATB2 P = 0.0419; CUX1 P = 0.430, two tailed Student’s t-test). **(e)** Representative caudal and rostral coronal sections of E17.5 control and poly(I:C) brains stained with DAPI. Yellow shading represents neocortical areas considered for thickness measurements. nCtx: neocortex, Th: thalamus, St: striatum, HC: hippocampus, Cp: caudate-putamen. **(f)** Quantification of cortical thickness measured at 30% and 70% distance from the dorsal midline and cortical hemispheric circumference (n = 5 control, n = 8 poly(I:C); caudal 30% P = 0.5252; caudal 70% P = 0.2481; rostral 30% P = 0.4459; rostral 70% P = 0.4583, two tailed Student’s t-test).

To investigate cell populations involved in the affected developmental processes, we analyzed astrocytes, oligodendrocytes and GABAergic interneuron cell populations. Concordant with RNA-seq results, the number of cells expressing GFAP, a protein found in radial glia and a subset of astrocytes (43), was increased in MIA offspring, mapping specifically to cells that seemed to emerge basally from the VZ **(Fig. 6a-b**, **Supplementary Fig. 8k)**. Since astrocytes also express SOX9 (44) and radial glia also express PAX6, which is decreased by MIA, this finding is consistent with the possibility that the increased SOX9+ cells outside the VZ, described earlier, may be indicative of newly differentiated astrocytes (**Fig. 4f)**. OLIG2, which is expressed in oligodendrocyte progenitors and mature cells as well as transiently in astrocyte progenitors (45), was also upregulated at E17.5 in the MIA group. OLIG2+ cells were increased in MIA brains compared to controls **(Fig. 6c-d, i**, **Supplementary Fig. 8k)**. The number of DLX2+ cells in the neocortex were comparable in the MIA and control groups (**Fig. 6e-f** **and** **6j**). In contrast, GAD immunoreactivity in this DLX2+ subpopulation, assessed by the pan-GABAergic marker GAD67+, was increased across the neocortex of MIA brains, and was not restricted to any specific zones of the neocortex (**Fig. 6g-h** **and 5k-m)**. Because GAD67 labels more mature cortical interneurons, this increase suggests alterations in interneuron development rather than total counts. While neuroanatomical characterization here was limited to E17.5 and interrogated a subset of genes and processes of interest, IHC findings correlate with transcriptional profiles, providing validation of MIA-associated DE in developing cortex.

**Figure 6.**
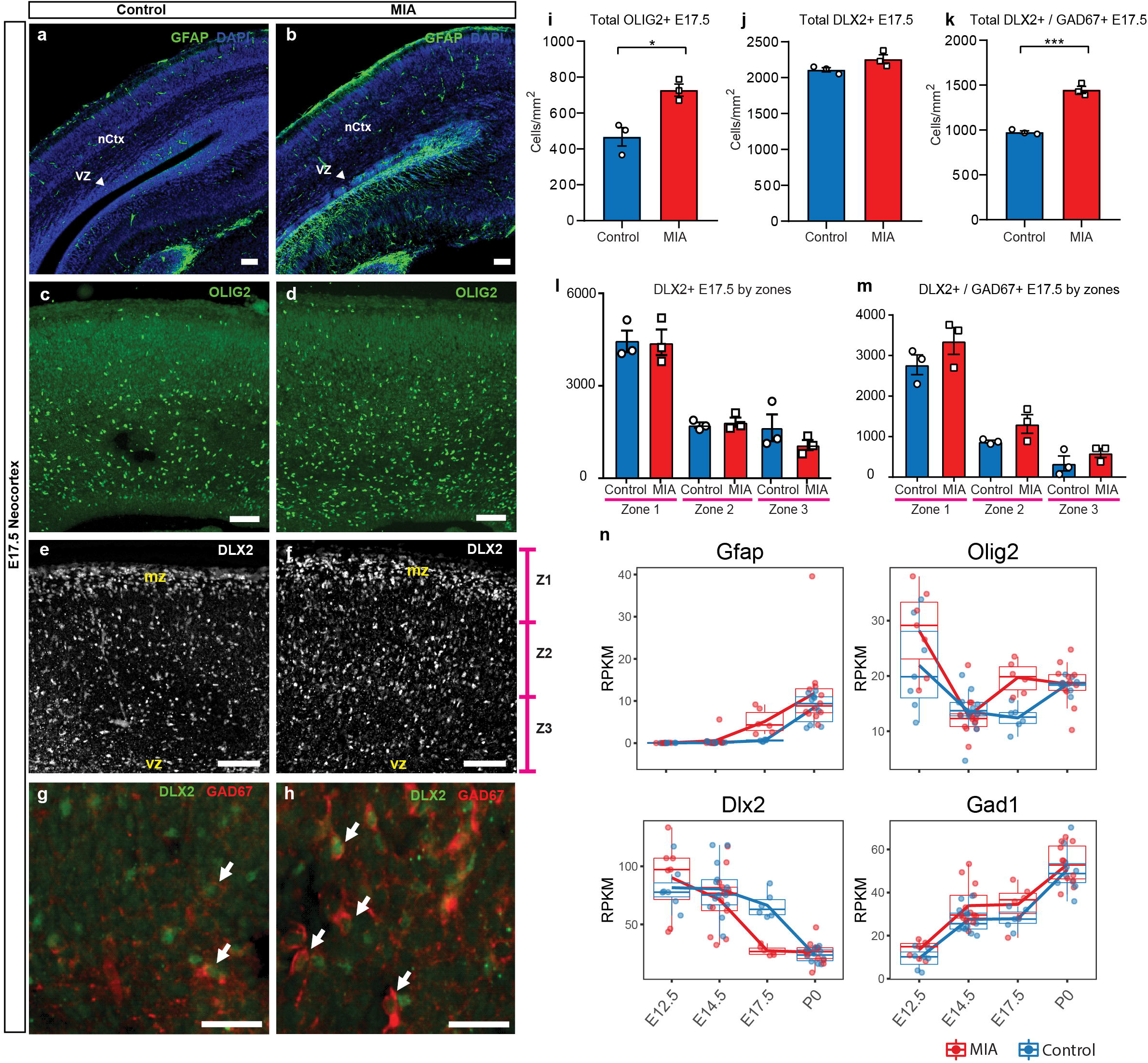
MIA alters development of neuronal and glial cell types in fetal offspring. **(a-h)** Coronal fetal brain sections from 2 independent E17.5 saline litters (n=4 control) and 3 independent MIA litters (n=6 MIA) were immunostained with markers to identify: radial glial cells and astrocytes using GFAP **(a, b)**, a sub-population of neuronal progenitors and oligodendrocytes using OLIG2 **(c, d),** and interneurons using the pan-GABAergic markers, DLX2 and GAD67. **(i, j)** Quantifications of the cell density of OLIG2 and DLX2 cells in the neocortex indicate an increase of OLIG2+ cells and no difference in DLX2+ cells (OLIG2 P = 0.0134 Student’s t-test). **(k)** Quantification of the cell density of double labeled DLX2/GAD67 cells in the neocortex shows an increase of DLX2+/GAD67+ cells. **(l, m)** Bin quantification analyses of DLX2+ and GAD67+ immunoreactive cells in zones 1-3 (Z1 to Z3). Arrows denote co-labeled cells. Scale bars in (a-f) = 100 μm and in (g-h) = 30 μm. Abbreviations: neocortex (nCtx); marginal zone (mz); ventricular zone (VZ). **(n)** RPKM trajectories of the cell identity markers tested in (a-k) across the developmental time points of the study.

## Discussion

Studies leveraging MIA models in mice have demonstrated the role of maternal immune signaling as a risk factor for cellular, anatomical, and behavioral outcomes in offspring. Nonetheless, a systems-level perspective of the early neurodevelopmental trajectory of pathology associated with MIA has been lacking. Our results offer a time-course resolved map of molecular targets aberrantly transcribed in fetal cerebral cortex following MIA. MIA-associated transcriptional signatures were robust, changed greatly across the surveyed time points, and revealed novel pathways and processes involved in pathology. Among MIA-associated DE genes, there was strong enrichment for ASD-relevant genes, providing evidence of the NDD-relevance of MIA transcriptional signatures. Both male and female samples were used, and we did not find evidence for strong sex-specific DE signatures. Elucidating the trajectory of transcriptional changes across embryonic cortical development following mid-gestational MIA represents a valuable step towards understanding the progression of MIA-induced pathology in the developing brain of offspring.

Our results suggest two waves of transcriptional pathology following MIA. First, we found initial induction of immune, metabolic, and angiogenesis gene expression modules specific to MIA exposure and largely restricted to the days following MIA. There is overlap in the acute induction signatures identified here and acute changes reported in other mid-gestational MIA models, including those using a range of injection gestational ages and rat models of LPS (8,11,12,16,46–48). Previous studies linked maternal immune challenge, prenatal brain hypoxia, and impacts on proliferation (49), neurogenesis, and cortical lamination (48). Thus, it is possible that the acute signatures identified here capture a generalized response in fetal brain across mid-gestational MIA models. Notably, induction of hypoxia, cellular metabolism and neuroimmune transcriptional signatures, particularly captured here by the Green module, persist at reduced levels at least through P0, and may indicate lasting perturbation to these pathways.

Of particular interest is the relationship between this acute response and downstream disruption to neurodevelopmental patterns of gene expression in the cortex. Identifying the early biological processes triggered by MIA are of critical interest for understanding how MIA initiates pathology. Our analysis identified potential drivers of these induced signaling pathways, including targets of substantial interest such as *Vegfa,* that link hypoxia, neurogenesis and cortical angiogenesis (50,51). *Vegfa* was upregulated in MIA-exposed animal at all time points, and was among the strongest DE genes at E12.5 and P0. Vegfa is critical for cortical development promoting neurogenesis and neurite outgrowth, as well as angiogeneisis (52–54). Upregulation of *Vegfa* is found in several NDDs and psychiatric disorders, including SZ and depression, and was also reported in a different MIA mouse model (55–62). Induction of angiogenesis in the developing cortex also independently causes a transition from proliferative state to neuronal maturation (54,63,64). Other genes of interest include *Pdk1* (pyruvate dehydrogenase kinase 1), which regulates glucose metabolism (65), Ldha (lactate dehydrogenase A), which facilitates glycolysis and participates in brain angiogenesis (66), *Bnip3* (BCL2 interacting protein 3), which is involved in cell death and mitochondrial autophagy (67), and *Il20rb* (interleukin 20 receptor subunit beta), a receptor subunit for pro-inflammatory interleukins IL-19, IL-20 and IL-24 (68). The early perturbed pathways observable within hours of MIA induction may be critical initiating steps leading to the altered general timing of fetal brain development described here. Our findings expand understanding of the molecular basis of these MIA-induced initial perturbations and reveal potentially key drivers of these changes.

The second component of the neurodevelopmental MIA signature defined here was perturbation to normal patterns of opposing proliferative and maturation neurodevelopmental modules accompanied by impacts on neuroanatomy and cortical lamination at E17.5. Initially, at six hours after MIA, there is a shift in expression of differentiation and neuronal maturation processes (e.g. synaptogenesis) to increased proliferative processes (e.g. positive regulation of cell cycle). This pattern parallels results described for the acute MIA response from other mouse and rat MIA models, where similar pathways were down-or up-regulated in fetal brain in the hours following MIA (69,70). Critically, our time course data shows that this initial response is reversed by E14.5, two days after MIA, with a peak in the reversed pattern of accelerated euronal maturation observed in MIA transcriptional signatures that was validated by neuroanatomical and protein-level changes in proliferative and maturing cell populations in the cortex at E17.5. Our time course design reveals that the transcriptional changes following MIA can be separated into discrete biological and temporal expression modules.

Our findings of disrupted cortical lamination following the acute MIA signature are consistent with, but not identical to previous studies showing cortical dysplasia associated with MIA-induced aberrant behavior in offspring (20,34). In those studies, dysplasia resolved at later postnatal stages but defects in lamination persisted into adulthood. Although we did not find evidence of aberrant cortical patches in the somatosensory cortex, secondary motor cortex or S1 and S1DZ, as previously reported, our results do show altered cortical lamination at E17.5. Thus, results from multiple studies using MIA models to date are consistent in showing a detrimental impact of MIA on cortical patterning. While significantly subtler in overall effects compared to earlier time points, we also see evidence for altered neuronal transcription and lasting immune activation at P0. While reduced in comparison to earlier timepoints, the changes in gene expression at P0 are consistent with deficits in synapse formation that require changes in expression of immune molecules in neurons from newborn MIA offspring (71).

Consistent with links between NDDs and developmental timing, atypical neurodevelopmental gene expression networks have been reported in a cortical neurodevelopmental model using autism-patient-derived induced pluripotent stem cells (54,72). Our findings suggest some aspects of the transition from neurogenesis to astrogenesis appear to occur at an earlier stage following MIA based on similar signatures of SOX9+ and GFAP+ cells reported in literature (73), and increased OLIG2+ cells and maturing GAD67+ DLX2+ interneurons in the cortex following MIA. These findings are additionally consistent with recent reports using poly(I:C) MIA models that identified perturbed GABAergic interneuron development (74), and maturation of GAD+ neuroblast after birth (75). Future work is needed to determine if and how MIA-associated dynamic transcriptional changes manifest postnatally and whether perturbation to embryonic development impacts cortical cytoarchitecture and function at later postnatal stages.

Altogether, our analyses indicate that MIA drives acute and lasting transcriptional changes that impact specific cell populations during development, disrupting normal developmental timing and causing increased numbers of astrocytes, oligodendrocytes, and altered molecular properties of GABAergic interneurons. The timing of MIA during gestation is well known to cause a range of phenotypes in offspring, and thus the findings here regarding mid-gestational MIA at E12.5 may not generalize to other times, especially for later gestational MIA after neurogenesis is largely complete. Further, the strong embryonic signatures we observe were generally consistent across sexes, raising the question of whether secondary insults or sex-specific protective effects contribute to some reports of sex differences in pathology following MIA (11,21,76). Nonetheless, the overlapping results between our study and other single time point studies of the acute MIA fetal brain transcriptional response suggest some consistency across bacterial and viral mimics, and rat and mouse models. Our study substantially expands previous work, revealing new molecular pathways and providing a temporal map of MIA-induced changes in across fetal neurodevelopment that will be relevant to understanding the pathology and informing future treatments of NDDs.

## Acknowledgments

This work was supported by NIH NIMH (project # 5R21MH116681-02), NARSAD Young Investigator Grant from Brain and Behavior Research Foundation and The UCD Clinical Translational Science Center. CPC is recipient of NIH NIMH Institutional National Research Service Award Autism Research Training Program fellowship (project # 2T32MH073124-16).

## Author Contributions

C.P.C., M.L.E. and K.C. are listed as joint first authors, as each led major components of the study; C.P.C., M.L.E., K.C., J.V.W., D.V., A.K.M. and A.S.N. designed the experiments and analyses; C.P.C., M.L.E., K.A., J.P.A., I.Z., E.J.K., E.C.C, J.M.S, K.F., T.S., D.V.L., J. B. G. and L.H. performed experiments; M.L.E, D.V.L and K.P. carried out mouse behavior; K.C. and A.S.N. performed and interpreted transcriptomic analysis; K.C., L.S.-F., and A.S.N. carried out WGCNA; C.P.C., K.A., and D.V. performed neuroanatomy analysis. C.P.C, M.L.E, K.C., D.V., A.K.M. and A.S.N drafted the manuscript. All authors contributed to manuscript revisions.

---

**Supplementary figure 1. Evaluation of females’ responsiveness to Poly(I:C) based on serum IL-6 levels.** Injection of poly(I:C) at E12.5 consistently elevates serum IL-6 in dams 4 hours following injection, compared with little response in dams injected with saline.

**Supplementary figure 2. Principal component analysis (PCA) of RNA-seq data. (a)** PCA plot with animal age represented as colors, and poly(I:C) / saline treatment indicated as circle / triangle shapes. **(b)** PCA plot with animal sex represented as colors, and poly(I:C) / saline treatment indicated as circle / triangle shapes.

**Supplementary figure 3. Males and females show concordant DE signatures following MIA. (a)** Numbers of upregulated and downregulated DE genes with p < 0.05 and FDR < 0.05, in the DE analysis lacking the sex covariate, are similar to the DE model including the sex covariate (Fig2c). **(b)** Numbers of upregulated and downregulated DE genes with p <0.05 and FDR < 0.05, in a DE analysis testing for differential expression between sexes identifies few DE genes, suggesting limited sex dimorphism between samples. MIA conditions and sequencing lanes were set as covariates for the DE model. **(c, d)** Numbers of upregulated and downregulated DE genes with p < 0.05 and FDR < 0.05, for the DE analysis performed on **(c)** males and **(b)** females demonstrate robust DE at E17.5 in both sexes.

**Supplementary figure 4. Sex-stratified comparison of DE genes and effect sizes. (a)** Correlation of the relative effect sizes (log_2_FC) for genes passing FDR < 0.05 identifies high degree of concordance between MIA responses between sexes. Genes passing FDR threshold in either or both sexes are color coded. **(b)** Venn diagrams illustrating overlap between DE gene sets passing FDR < 0.05 at E14.5 and E17.5, in DE models containing samples from both sexes, males, or females. The DE model comprising individuals of both sexes includes most of the DE signature of each sex. Female DE analysis identifies limited and unique DE signature. **(c)** Examples of E14.5 and E17.5 genes significant in females but not males. At E14.5 genes were selected to pass FDR < 0.05 and logFC > 0.5 or logFC < −0.5 in females, but not logFC > 0.5 or logFC < −0.5 in males. The gene-set selection process was analogous for E17.5 with the addition of FDR < 0.05 condition in males. After gene set filtering, twelve genes with the lowest FDR in females were plotted.

**Supplementary figure 5. Summary of WGCNA module eigengene expression, association with experimental variables, and overlap with DE genes. (a)** Eigengene trajectories across neurodevelopment of the ten original co-expression modules. **(b)** Heatmap of statistical association between gene expression modules and experimental traits; age, condition (saline vs poly(I:C)), and sex. Based on the similarities of temporal gene expression trajectories six of the original ten modules were grouped into two, BrRePi: Brown, Red and Pink; YeMaBl: Yellow, Magenta and Black. Numerical values represent signed Pearson’s correlation coefficients, and, in brackets, Student asymptotic P values. Green color represents negative, red color represents positive correlation. Color intensity signifies strength of the correlation. Blue and Turquoise modules are strongly associated with age; Green and Grey modules are significantly associated with the condition. **(c)** DE (p <0.05) gene set enrichment analysis in WGCNA modules across the developmental time points of the study. Values representing the number of upregulated and downregulated genes are plotted above and below 0, respectively. Asterisks indicate statistically significant enrichment of DE genes (P < 0.05) in a module (p <0.05; hypergeometric test).

**Supplementary figure 6. Heatmaps of GO BP enrichment analysis of upregulated and downregulated differentially expressed genes in WGCNA modules. (a)** E12.5 upregulated DE genes (FDR < 0.05). **(b)** E12.5 downregulated DE genes (FDR < 0.05). **(c)** E14.5 upregulated DE genes (FDR < 0.05). **(d)** E14.5 downregulated DE genes (FDR < 0.05). **(e)** E17.5 upregulated DE genes (FDR < 0.05). **(f)** E17.5 downregulated DE genes (FDR < 0.05). **(g)** P0 upregulated DE genes (FDR < 0.05). **(h)** P0 downregulated DE genes (FDR < 0.05). Only modules with more than 5 DE genes (FDR < 0.05) are shown in the heatmaps.

**Supplementary figure 7. Sex-stratified RPKM gene trajectories of genes presented in the manuscript showing concordant effects between males and females. (a)** Vegfa, Flt1, Pdk1, Ldha, Bnip3, Il20rb. **(b)** Mki67, Eomes, Pax6, Sox9, Tbr1, Bcl11b, Satb2, Cux1. **(c)** Gfap, Dlx2, Olig2, Gad1.

**Supplementary figure 8. Raw Western blot and RNA-seq supporting data to proliferation, lamination, and cell-specificity analyses. (a)** Western blot demonstrates reduced PAX6 protein level in cortical extracts from poly(I:C) treated animals at E17.5, **(b)** but not at P0. **(c)** Quantified PAX6 protein levels from Western blot analysis at P0 (n = 6 for each control and poly(I:C) groups; P = 0.766, two tailed Student’s t-test). **(d-g)** RNA-seq RPKM expression trajectories throughout development of **(d)** Ki67, **(e)** Tbr2, **(f)** Pax6, and **(g)** Sox9. **(h)** Individual Western blots detecting TBR1, CTIP2, SATB2 and CUX1 from whole cortical extracts at E17.5 and **(i)** CTIP2 at P0. **(j)** E17.5 RNA-seq RPKM expression levels of Pax6, Neurog2, Notch1, and **(k)** Gfap and Olig2.

