## Supplementary methods for "A temporal map of maternal immune activation-induced changes reveals a shift in neurodevelopmental timing and perturbed cortical development in mice"

**Immunohistochemistry staining**

Fetal brains were dissected and fixed with 4% paraformaldehyde/PBS solution overnight at 4°C. Tissues were then equilibrated in approximately 15 mL of 30% sucrose/PBS solution until they sank to the bottom of a conical tube. Equilibrated brains were embedded in Optimum Cutting Temperature (OCT) compound (Tissue-Tek, Torrance, CA) and frozen on dry-ice. OCT embedded brain blocks were cryo-sectioned on a coronal plane (30 µm). Immunostaining was performed in free-floating sections with agitation. Sections were washed five times in PBST (PBS with 0.05% Triton X-100, 5 min each) and antigen retrieval was performed using 1x Citrate buffer pH6.0 antigen retriever solution (cat# C9999; Millipore-Sigma, Burlington, MA ), at 60°C for 1 hour. Sections were washed 5 times in PBST (5 min each), permeabilized in PBS containing 0.5% Triton for 20 min and blocked for 1h at room temperature in 5% milk/PBST. Primary antibodies were incubated overnight at 4°C with orbital agitation (40-50 rpm). All antibodies used for this study were validated and their use widely reported. The following primary antibodies were used: anti-PAX6 (1:250; cat #PRB-278P-100; Covance, Princeton NJ.), anti-KI67 (1:500; cat#12202; Cell Signaling, Danvers, MA), anti-SOX9 (1:500 cat# AF3075, R&D systems, Minneapolis, MN), anti-PH3 (1:500 cat# 9701, Cell Signaling, Danvers, MA), anti-TBR2 (1:500; cat#14-4875-82; Thermo Fisher Scientific, Waltham, MA.), anti-TBR1 (1:500; cat#ab31940; Abcam, Cambridge, MA), anti-CTIP2 (1:250; cat# ab18465; Abcam, Cambridge, MA), anti-CUX1 (1:200; cat#A2213; Abclonal, Woburn, MA.), anti-SATB2 (1:500; cat#ab51502; Abcam, Cambridge, MA.), anti-DLX2 (1:200; generous gift from John Rubenstein, UCSF), anti-GFAP (1:250, cat#Z0334; Agilent Dako, Santa Clara, CA.), anti-OLIG2 (1:500, cat#AB9610, MilliporeSigma, Burlington, MA). After primary antibody incubation, free-floating sections were washed five times in PBST (5 min each). Species-specific fluorophores-conjugated IgG (1:1000; Invitrogen-Thermo Fisher Scientific, Waltham, MA) were used as secondary antibodies (45 min, RT). 40,6-Diamidino-2- phenylindole (DAPI) (1:10000; Millipore-Sigma, Burlington, MA) was used for nuclear staining (20 min, RT).

**Immunoblotting**

Isolated forebrain from E17.5 embryos (male and female littermates) from at least 3 litters per condition were dissected in HBSS, immediately frozen on dry ice, and stored at -80°C until processed. Samples were lysed in 50 mM Tris HCl, pH 8, 140 nM NaCl, 1 mM EDTA, 10% glycerol, 0.5% NP40 and 0.25% Triton with protease inhibitor cocktail (Roche). After sonication, samples were spun down and the supernatant was used for a BCA Bradford assay using the Spectramax 190 plate reader to assess protein concentration using a standard curve. 12 or 18 μg of protein were run on a 10 or 12% Tris acetate gel using the Mini-PROTEAN system (BioRad). Membranes were blocked in Odyssey blocking buffer (TBS; LiCor) and probed with the indicated primary antibodies overnight at 4°C. Resolved proteins were visualized in two channels using fluorescent secondary antibodies at 680 and 800nm on an Odyssey Clx infrared imaging system (LiCor). Specific band intensities for all detected antibodies were quantified using the manufacturer’s software ImageSoft (LiCor) and normalized to GAPDH loading control. Antibodies used were anti-PAX6 (1:250; cat #PRB-278P-100; Covance, Princeton NJ.), anti-TBR1 (1:500; cat#ab31940; Abcam, Cambridge, MA), anti-CTIP2 (1:250; cat# ab18465; Abcam, Cambridge, MA), anti-CUX1 (1:200; cat#A2213; Abclonal, Woburn, MA.) and anti-SATB2 (1:500; cat#ab51502; Abcam, Cambridge, MA.).

**Bioinformatics analysis**

Bioinformatic analysis was performed using R programming language version 3.5.1 (1) run in RStudio integrated development environment version 1.2.1269 (2). Plots were generated using ggplot2 R package version 3.1.0 (3). Heatmaps were generated using pheatmap R package 1.0.10 (4).

**Differential expression analysis**

Raw count data for all samples were used as input along with sample information for differential expression analysis using edgeR (5). Genes with minimum log_2_ reads per kilobase per million (RPKM) expression of -2 in at least two samples were included for analysis, resulting in a final set of 17,195 genes for differential testing. For time point differential expression analysis genes with count per million (CPM) > 0.1 in at least 2 samples were considered, resulting in E12.5, 16,396; E14.5, 16,658; E17.5, 16,229; P0, 16,613 genes in differential expression analyses. Principal component analysis indicated that the strongest driver of variance across samples was developmental age. Tagwise dispersion estimates were generated, and differential expression analysis was performed with edgeR using a generalized linear model including sex, sequencing run factor, and devel­opmental stage as the variable for testing. Stage-specific differential expression testing was also performed. Normalized expression levels were generated using the edgeR rpkm function. Normalized log_2_ RPKM values were used for plotting of summary heatmaps and of expression data for individual genes.

**SFARI gene set enrichment analysis**

The autism risk gene-set was downloaded from https://gene.sfari.org/ on 02-26-2019 and is included in this manuscript as **Supplementary Table 19**. High confidence risk genes, annotated as “gene-score” 1 and 2 were selected, and their orthologs were found using getLDS function from the biomaRt R package (6,7). Overlap of up- and downregulated DE genes with SFARI ortholog genes was calculated and plotted using a custom R script. Statistical significance of overlap was tested with hypergeometric test using the following R script:

*sum(dhyper(t:b, a, n - a, b))*

*t* = number of overlapping DE genes with SFARI gene orthologs with gene_score 1 or 2

*b* = number of SFARI gene orthologs with gene_score 1 or 2

*a* = number of DE genes at a developmental stage

*n* = number of genes at a developmental stage

*sum* returns the sum of all the values present in its arguments.

*dhyper* calculates density for the hypergeometric distribution.

**WGCNA.**

We used the WGCNA R package, version 1.66 (8,9) to construct signed co-expression networks using the entire dataset containing 24,015 genes. After the network construction, the gene set was filtered for minimal gene expression at an RPKM value of 0.25 or higher in at least two sample, resulting in a dataset consisting 17,195 genes. A correlation matrix using the biweight midcorrelation between all genes was computed for all relevant samples. The soft thresholding power was estimated and used to derive an adjacency matrix exhibiting approximate scale-free topology (*R*^2^ > 0.85). The adjacency matrix was transformed to a topological overlap matrix (TOM). The matrix 1-TOM was used as the input to calculate co-expression modules using hierarchical clustering.

Modules were branches of the hierarchical cluster tree base, with minimum module size set to 10 genes. In addition, Pearson’s correlation coefficients were used to calculate correlation between sample traits (e.g., sex, treatment) and modules. The expression profile of a given module was summarized by the module eigengene (ME). Modules with highly correlated MEs (correlation > 0.80) were merged together. The module connectivity (kME) of each gene was calcu­lated by correlating the gene expression profile with module eigengenes. The module connectivity (kME) of each gene was calcu­lated by correlating the gene expression profile with module eigengenes. Genes with no network correla­tion were placed into the module Grey. Following manual data inspection further highly correlated modules were merged: BrRePi= Brown + Red + Pink, YeMaBl = Yellow + Magenta + Black.

**Gene Ontology enrichment analysis.**

Mouse Gene Ontology (GO) data was downloaded from Bioconductor (org.Mm.eg.db). We used the TopGO R package version 2.34.0 (10) to test for enrichment of GO terms. For the analysis presented here, we restricted our testing to GO Biological Process annotations and required a minimal node size (number of genes annotated to GO terms) of 20. We used the internal ‘weight01’ testing framework and the Fisher test, a strategy recommended for gene set analysis that generally accounts for multiple testing comparisons. For GO BP analysis, we reported terms with p-value < 0.05. For all enrichment analysis, the test set of DE genes was compared against the background set of genes expressed in our study based on minimum read-count cutoffs described above. Heatmaps showing positive log_2_(expected/observed) values were plot­ted for GO terms of interest.

**Protein–protein interaction**

Protein–protein interaction enrichment and network generation for E12.5 DE (FDR < 0.05) gene sets was performed using STRING (11), version 11, considering only experimentally and text mining interactions, with at least medium interaction confidence score. Disconnected nodes were removed from the network.

**Supplementary Methods References**

1. R Core Development Team (2015): R: a language and environment for statistical computing, 3.2.1. *Document Freely Available on the Internet at: Http://Www. r-Project. Org*. Vienna, Austria: R Foundation for Statistical Computing. https://doi.org/10.1017/CBO9781107415324.004

2. Team R (2018): RStudio: Integrated Development Environment for R. Boston. Boston, MA.

10. Alexa A, Rahnenfuhrer J (2019): topGO: Enrichment Analysis for Gene Ontology. R package version 2.38.1.

11. Szklarczyk D, Gable AL, Lyon D, Junge A, Wyder S, Huerta-Cepas J, *et al.* (2018): STRING v11: protein-protein association networks with increased coverage, supporting functional discovery in genome-wide experimental datasets. *Nucleic Acids Res* 47: 607–613.
