## Supplementary figures and images for "A temporal map of maternal immune activation-induced changes reveals a shift in neurodevelopmental timing and perturbed cortical development in mice"

### Sup Fig 1

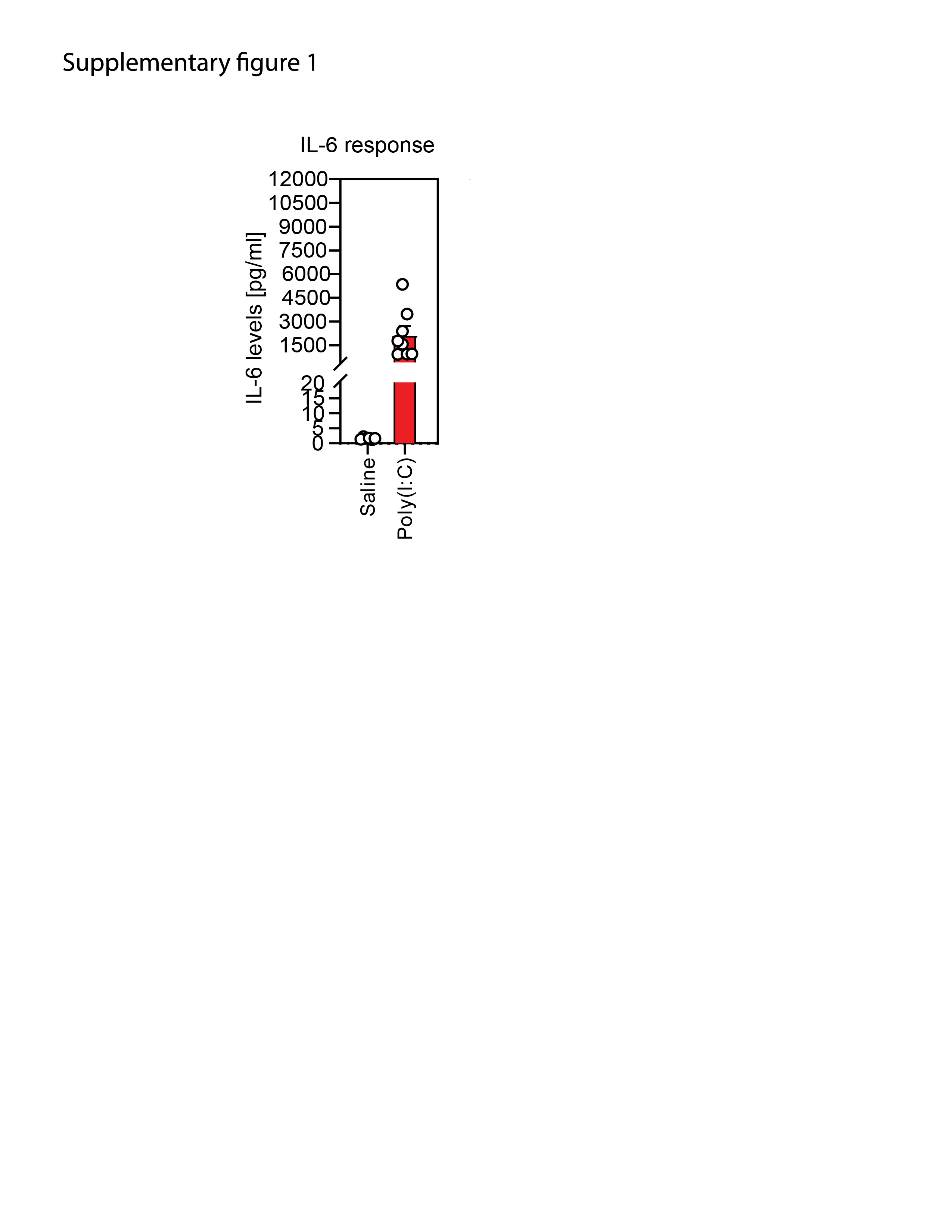

### Sup Fig 2

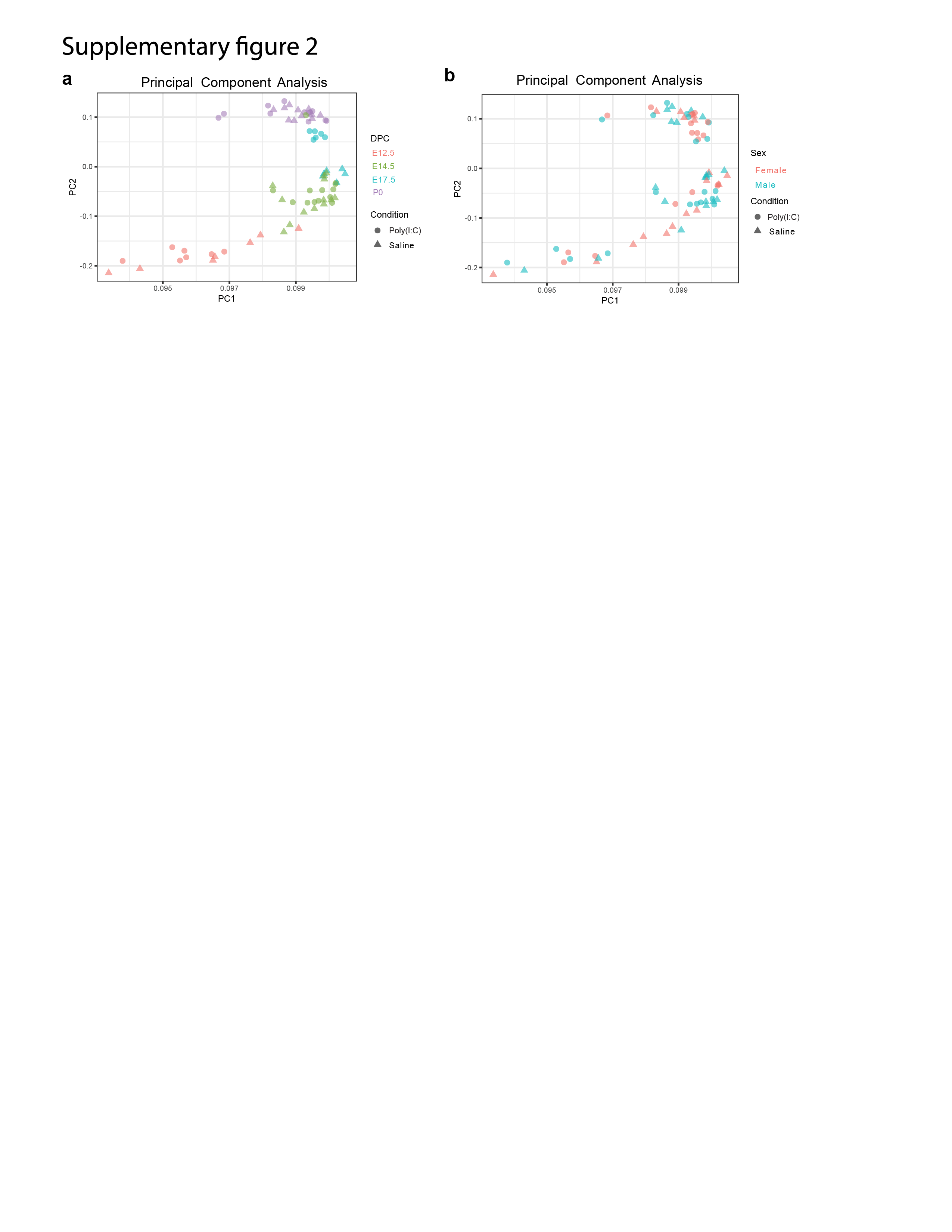

### Sup Fig 3

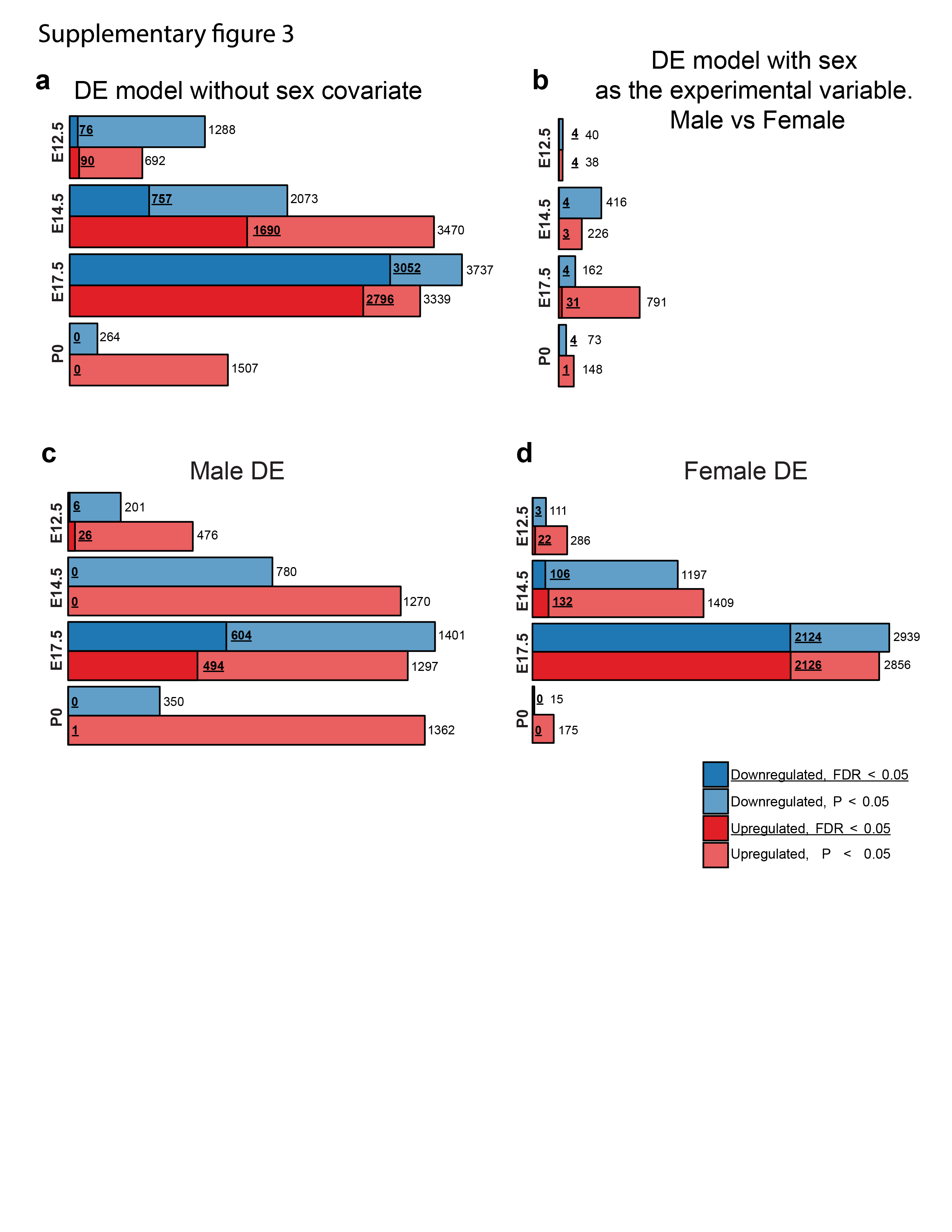

### Sup Fig 4

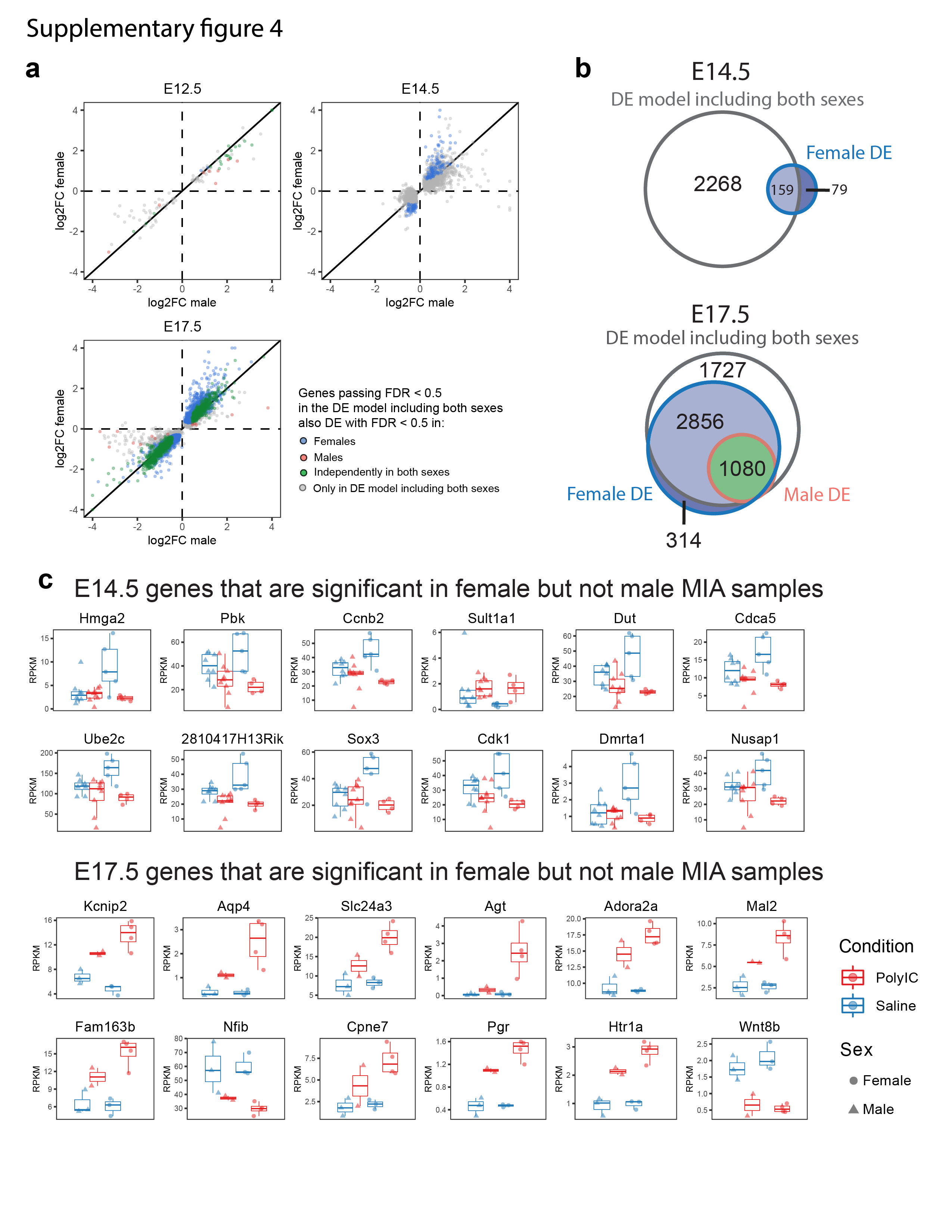

### Sup Fig 5

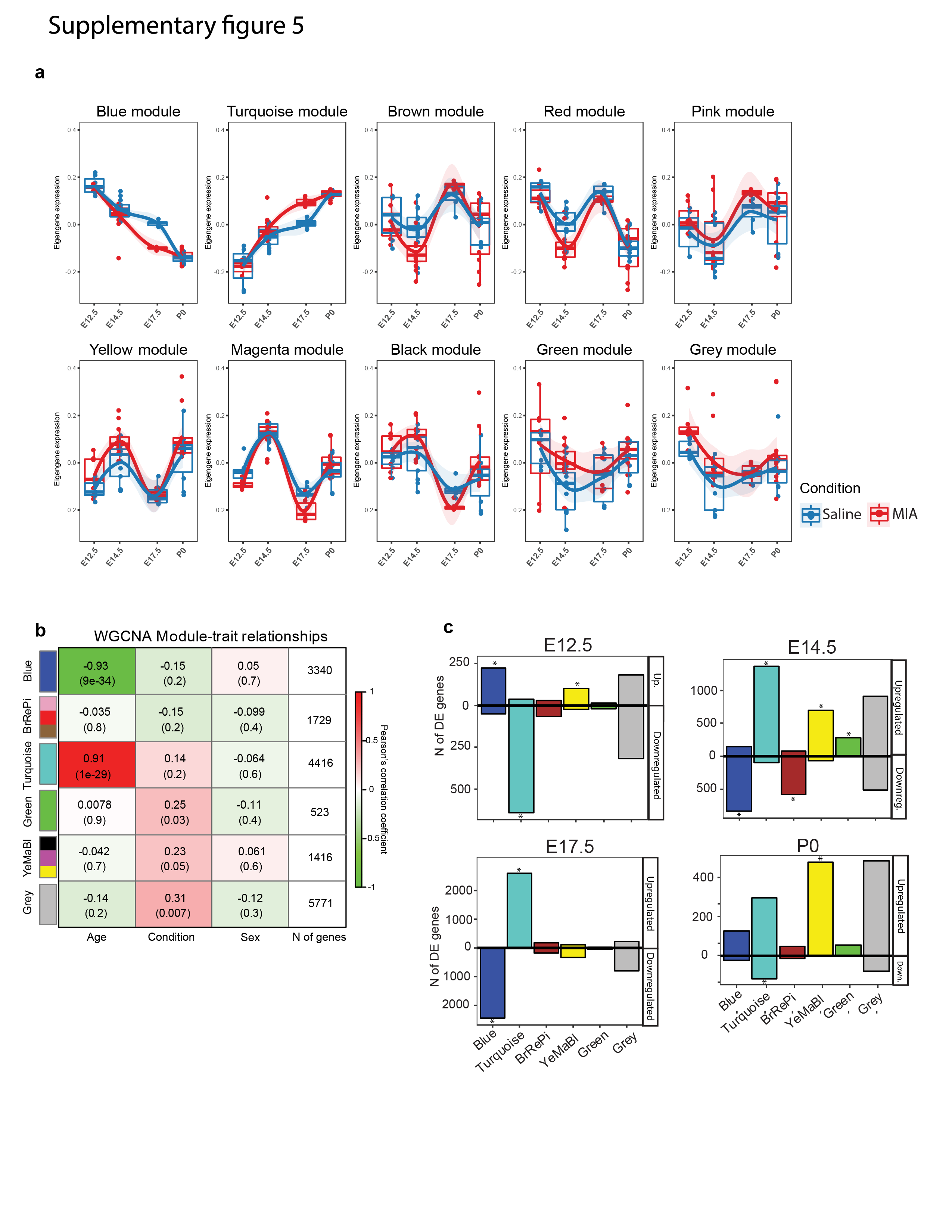

### Sup Fig 6

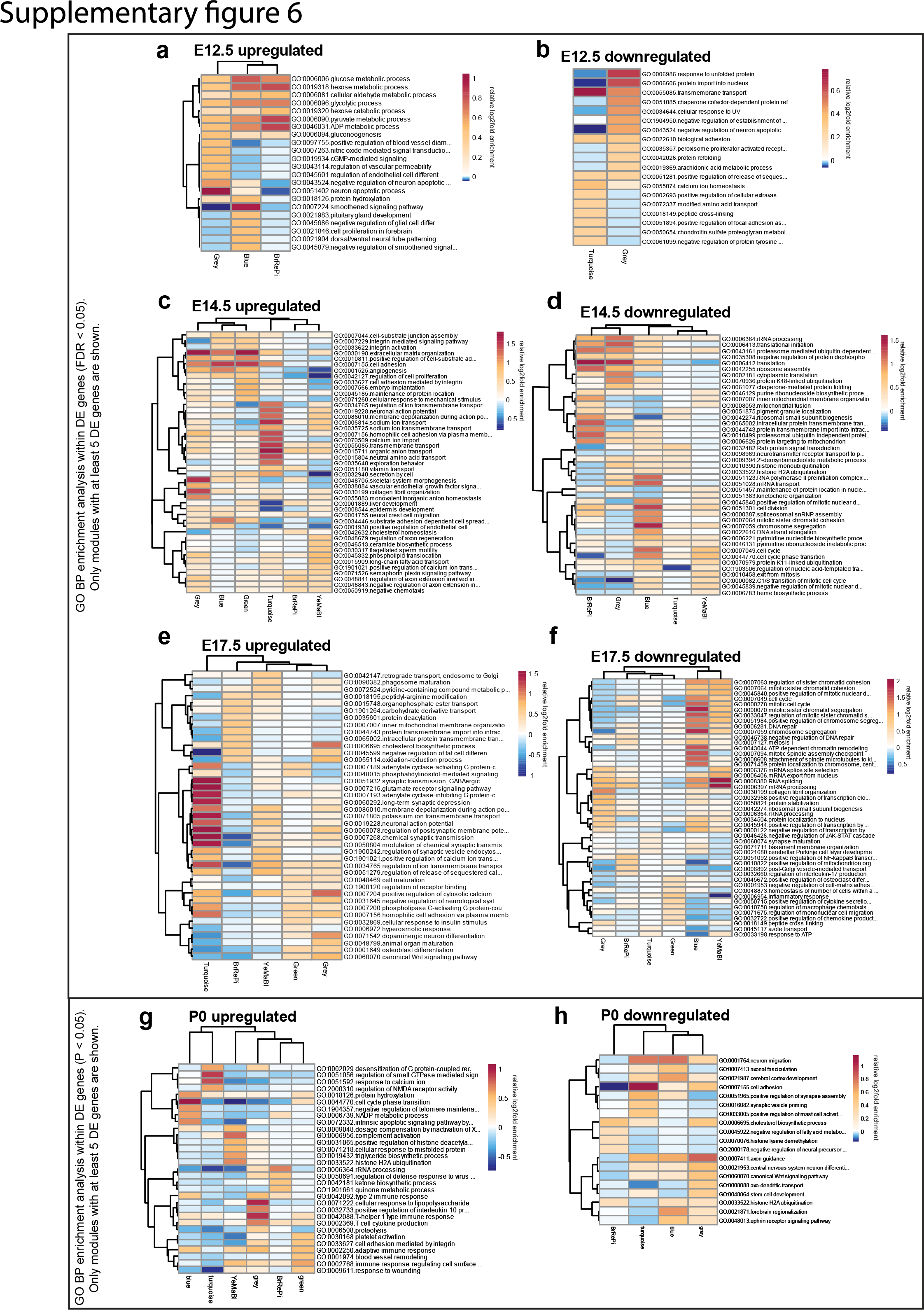

### Sup Fig 7

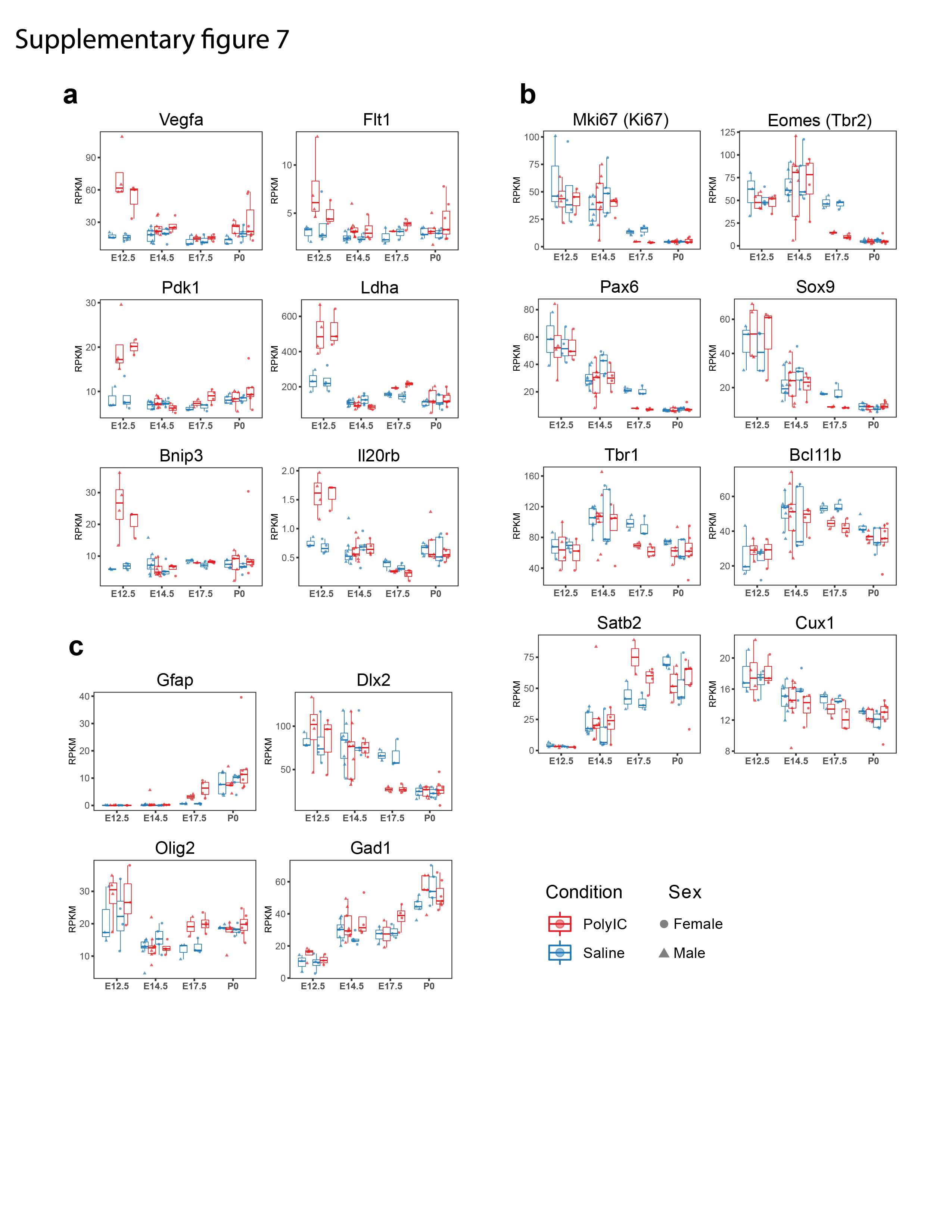

### Sup Fig 8

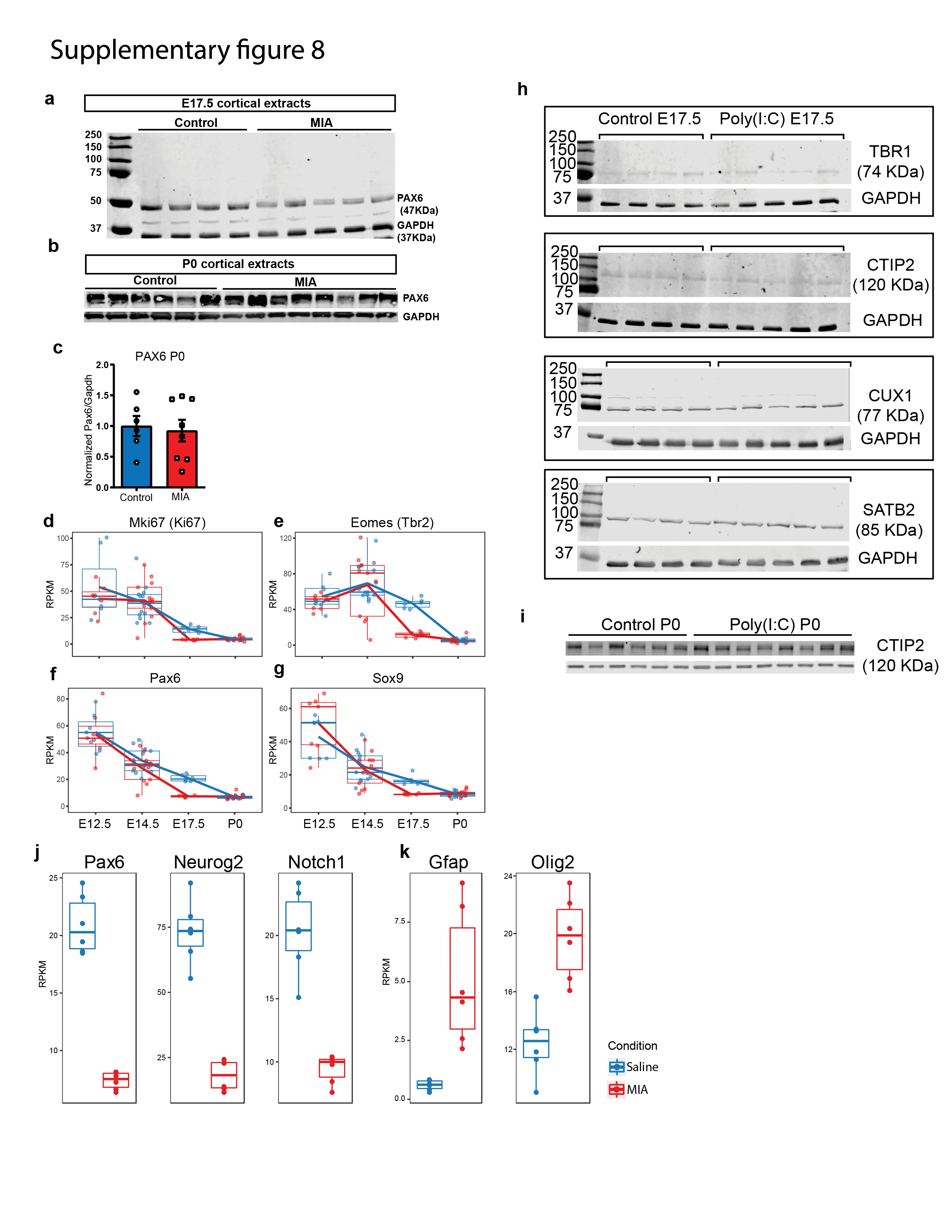
